# An Automated Wireless Seesaw System Enabling Spatial Separation of Action and Reward in Group-Housed Marmosets

**DOI:** 10.64898/2026.06.02.728149

**Authors:** J Cabrera-Moreno, J. M. Burkart, K. R. Brügger

## Abstract

Cooperation in social species is shaped by ongoing social relationships, partner choice, and group interactions, demanding experimental systems that preserve the social context in which these behaviors unfold. Here we introduce the e-Seesaw, a wireless system for automated liquid reward delivery designed to support home-enclosure experiments on reward access distribution in common marmosets (*Callithrix jacchus*) with minimal human intervention. The apparatus combines a modular peristaltic pump with a Bluetooth-controlled trigger, allowing spatial separation between the site of action and the site of reward delivery while preserving group housing. We provide detailed design files, software, and assembly instructions to support reproduction and adaptation. In a proof-of-concept deployment across seven families, animals readily engaged with the device, producing a median of ~87 trigger activations per session. Engagement was concentrated early within sessions and remained largely stable across repeated deployments, including under increased action-reward separation. These results established the e-Seesaw as a flexible and reproducible platform for automated reward-delivery experiments in animals tested within their social groups, while reducing human involvement and avoiding fixed dyadic testing.

## Introduction

Understanding the evolution of human cooperation and the neural and cognitive processes that support it remains a major question across evolutionary anthropology, comparative psychology and neuroscience (*1*–*7*). Because cooperative behavior unfolds through interactions with partners, and depends on social relationships (e.g., (*8, 9*)), shared outcomes (e.g., *(10, 11)*), and conflicting interests (e.g., (*12, 13*)), it is important to study it under conditions that preserve the interaction structure in which it naturally occurs while minimizing human disruption. Addressing this challenge therefore requires conceptual frameworks that guide comparative research (*14*) and experimental paradigms that permit controlled assessment of cooperative behavior in socially relevant contexts (*15*).

Experimental paradigms studying cooperation and prosocial behavior typically seek to place reward structures under experimental control, quantify individual contributions to cooperative outcomes, and assess how animals respond to the presence or behavior of partners. This general logic is captured by a range of tasks, including the prosocial choice test (PCT) (*16*), in which paired animals choose between reward distributions that either favor both the actor and the partner (mutualistic choice), only the partner (prosocial choice), or only the actor (selfish choice) (*17*–*22*). A related extension of this approach is the “group service paradigm”, which retains a similar logic while moving beyond dyadic interactions to group settings and alleviates many of the difficulties of the PCT with reward pay-off distributions (*23*–*34*).

Although these paradigms have shed light on a variety of questions regarding the evolution of cooperation and prosocial behavior, their implementation often relies on testing situations that require extensive human intervention. In practice, this can slow data collection through repeated resetting of the apparatuses’ position, reloading rewards between trials or initiating trials (*15*), and continued reliance on manual video coding, ultimately limiting the dataset size and statistical power especially in the face of high interindividual variability in prosocial propensities. At the same time, such procedures can reshape the very social conditions under which cooperation is expressed, for example by requiring temporary separation from their social group or imposing specific dyads to work together (e.g., (*20, 22, 35*–*37*)). This is especially relevant because cooperative and prosocial responses often depend on who the partner is in terms of rank, kinship or sex and on whether individuals can choose their interaction partners (*27, 33, 33, 35, 38*–*41*). More broadly, in nonhuman primates, this concern extends to the experimenter itself, which may become part of the social situation rather than a neutral background factor (*42*). Finally, conventional setups often constrain the spatial positioning of apparatuses, making it difficult to integrate experiments flexibly into home enclosures or to manipulate the spatial relationship between action and reward delivery.

Recognizing the limitations of traditional testing paradigms, researchers have increasingly developed automated systems that enable animals to participate voluntarily in cognitive experiments within their home enclosures (*43*–*47*). Such approaches can improve animal welfare due to reduced stress from separation of the family group, increase experimental throughput, and allow for more naturalistic social contexts during testing as well as improve ecological validity by removing the need for the presence of a human experimenter during testing. Automated paradigms have also been developed more specifically to test cooperation in various species, including touchscreen-based prosocial choice tasks (*48, 49*), liquid-based group service task variants (*34, 38, 50*).

However, most existing solutions rely on fixed interaction stations, in which the site of action and reward delivery are co-located, or on task-specific apparatuses that are difficult to reposition within the enclosure due to their size or their materials. To address these constraints, we developed the e-Seesaw (*26, 29*), an open-source, wireless seesaw-based system for automated liquid reward delivery. The system couples a Bluetooth-controlled trigger to a modular peristaltic pump, enabling spatial separation between the location of animal interaction and the site of reward delivery while keeping the liquid reward and electronics protected from direct contact with the animals. Its compact, enclosure-compatible design supports easy rearrangement of trigger and reward placement across experimental layouts without changes to the housing infrastructure. By making complete 3D printable design files, software, and assembly instructions openly available, the e-Seesaw offers a reproducible foundation for home-enclosure experiments requiring spatially displaced reward delivery.

Common marmosets (*Callithrix jacchus*) are a suitable model for deploying such technology. As cooperative breeders living in stable family groups, they show high levels of social tolerance and routinely interact around feeding contexts, including co-feeding and, in some settings, food transfers (*20, 51*–*53*). Marmosets are also increasingly used in behavioral biology and neuroscience due to their tractable size, suitability for more naturalistic housing, and growing availability of tools for recording and manipulating neural activity (*54*).

Here, we assess the deployment feasibility of the e-Seesaw under routine home-enclosure conditions across seven marmoset families (4–5 individuals), including tests with increased spatial separation between the trigger and reward module. We quantify usage dynamics across families and sessions to determine whether group-housed marmosets engage with the device reliably over repeated daily deployments. In these proof-of-concept tests, animals engaged readily with the system across sessions, producing a median of ~87 trigger activations per session (median session duration = 100 min), with half of all activations occurring within the first ~23% of session duration. Engagement trajectories across families and repeated sessions were largely stable, with two families showing a robust increase in early-session engagement over time.

## Results

We evaluated the e-Seesaw system (Fig. 1A) under routine home-enclosure conditions across 31 adult common marmosets of either sex and housed in groups of 4–5 members, all of which had prior experience with a functionally similar seesaw mechanism (see Methods). Animals had ad libitum access to the device during routine deployment, so session duration was not experimentally fixed but reflected the daily husbandry routine. Unless otherwise noted, the separation between the interaction site and reward outlet was 20 cm. To assess deployment feasibility and characterize device use across days, we first summarize the number of sessions and interaction incidence across families, then characterize how triggers were distributed within sessions, and evaluate whether early-session engagement changed systematically across repeated sessions within each family. Finally we report a subset of deployments at larger action–reward separation, 100 cm.

**Figure 1.**
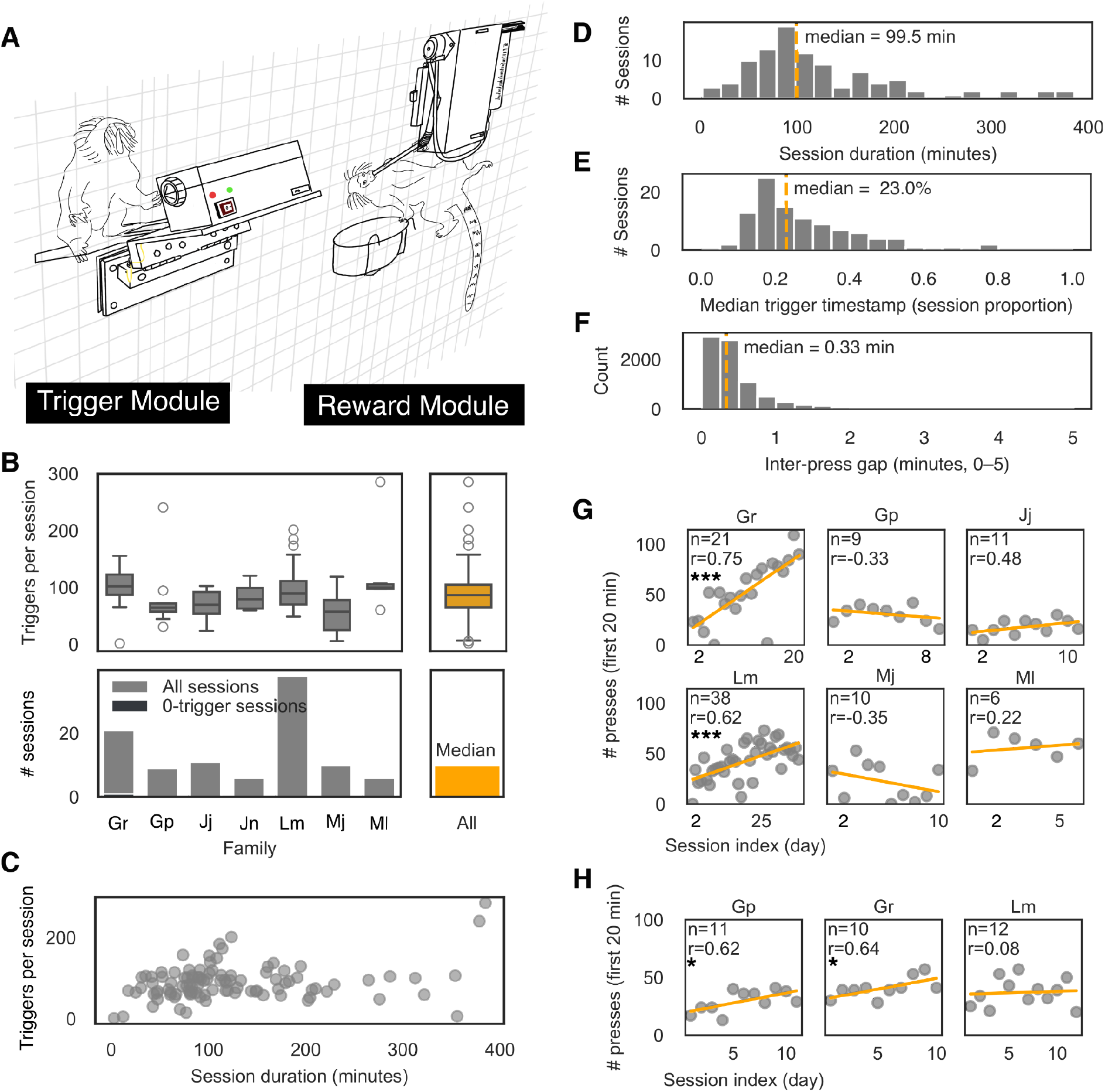
(**A**) Schematic representation of two marmosets using the e-Seesaw in their home enclosure. (**B-upper**) Session-level engagement across families, shown as the distribution of triggers per session for each family (boxplots, each point represents one session). (**B-lower**) Sampling coverage across families: number of sessions per family with overlay indicating zero-trigger sessions.(**C**) Relationship between session duration and total triggers per session (Pearson correlation r = 0.23). (**D**) Distribution of session durations (minutes), computed as elapsed time between first and last recorded event within a session. The dashed line denotes the median. (**E**) Distribution across sessions of the median trigger timestamp expressed as a fraction of session duration. The dashed line indicates the median across sessions. (**F**) Distribution of inter-trigger gaps within sessions (minutes, truncated display range 0-5 minutes). The dashed line denotes the median gap. (**G**) Early-session engagement trajectories: triggers occurring within the first 20 minutes of each session plotted across successive sessions for each family with fitted linear trend lines. Asterisks indicate whether the fitted slope differs from zero: * p < 0.05, ** p < 0.01, *** p < 0.001. (**H**) Similarly as in G but in deployments with a 100-cm separation between the trigger and reward module, shown for a subset of 3 families.

### Session-level engagement and sampling coverage

Session-level engagement, defined as the number of triggers per session, showed moderate variability across families (Fig. 1B-top). Across sessions, the median number of triggers per session was 87 (IQR = 65.8–105.3), with values ranging from 2 to 286 triggers. A total of 101 sessions were analyzed, with families contributing 6 to 38 sessions each (median per family = 10 sessions, Fig. 1B-bottom). Sessions with no triggers were rare (median per family = 0 zero-trigger sessions, range 0–1, Fig. 1B-bottom), indicating near-universal interaction when the device was available.

### Session duration is heterogeneous across days

Session duration, defined as the elapsed time between the first and last recorded trigger within a day, varied widely across sessions (Fig. 1C). Across sessions with non-zero duration (n=100), durations ranged from 2.6 min to 384.7 min (Fig. 1C). The median session duration was 99.5 min, with an interquartile range of 73.1–159.1 min (Fig. 1D). The coefficient of variation (CV, SD/mean) for session duration was 0.67, indicating substantial heterogeneity in sampling across days.

### Engagement within sessions is strongly front-loaded

To characterize engagement within individual sessions, trigger timestamps were expressed relative to each session’s total duration, allowing comparison across sessions despite substantial variation in session length (Fig. 1E). This normalization was relevant because device access was granted ad libitum rather than within a fixed testing interval. Despite wide variation in session duration (Fig. 1C), engagement was temporally concentrated early in sessions (Fig. 1E). The median trigger timestamp within a session occurred at 23% of session duration and at 21.7 min after session onset in absolute time, indicating strong front-loading of interaction within individual sessions. Consistent with sustained interaction during active periods, the median inter-trigger gap was 0.35 min (21 s), with 90% of gaps < 2 min and 95% < 4.6 min (Fig. 1F), indicating continuous engagement when animals interacted with the device. This pronounced front-loading suggests that later portions of long sessions contribute relatively little additional engagement and that total session duration is a poor proxy for opportunity to engage.

Consistent with the within-session timing results, session duration explained only a small fraction of the variance in trigger counts. Across sessions, the correlation between total session duration and number of triggers was r = 0.23, corresponding to ~5% of variance explained. This weak relationship indicates that longer sessions do not simply scale engagement proportionally and supports treating engagement as a session-level behavioral choice rather than a time-normalized rate.

### Engagement persists across sessions

To quantify day-to-day changes in engagement while limiting confounding by variable session duration, we defined engagement as the number of trigger activations occurring within the first 20 minutes of each session. This fixed window standardizes the metric across sessions and was selected based on the observed front-loading of presses within sessions. Sessions were ordered chronologically within each family, and linear trends were estimated across sessions. Under a combined criterion requiring both statistical support (95% confidence interval excluding zero) and a practically meaningful effect size (≥±5% change per session relative to the family median), five of seven families were classified as stable. Two families showed a robust increase in early-session engagement over time, and these were also among the families sampled across the largest number of sessions (Table 1, Fig. 1G). Thus, repeated exposure did not lead to a detectable decline in engagement, although longer deployments may reveal increasing engagement trajectories in additional families.

**Table 1.** For each family, the table reports, the group size, the number of sessions analyzed, the median number of triggers within the first 20 minutes of a session, the estimated linear slope of early-session triggers per session (with 95% confidence interval), the relative slope (slope normalized by the family median), the Pearson correlation coefficient, the slope p-value and FDR-corrected p-value.

| family | group size | n_sessions | median_presses | slope_per_session | slope_ci_lo | slope_ci_hi | relative_slope | pearson_r | p_slope | p_slope_fdr |
| --- | --- | --- | --- | --- | --- | --- | --- | --- | --- | --- |
| Gp | 4 | 9 | 34.0 | -1.033 | -3.683 | 1.617 | -0.03 | -329 | 387 | 542 |
| Mj | 4 | 10 | 21.0 | -2.176 | -6.971 | 2.62 | -104 | -347 | 326 | 542 |
| Jj | 4 | 11 | 16.0 | 1.064 | -388 | 2.515 | 66 | 484 | 132 | 307 |
| Gr | 4 | 21 | 52.0 | 3.59 | 2.083 | 5.096 | 69 | 753 | 0.0 | 0.0 |
| Jn | 5 | 6 | 38.0 | -2.343 | -10.498 | 5.812 | -62 | -0.37 | 0.47 | 548 |
| MI | 5 | 6 | 59.5 | 1.629 | -8.296 | 11.553 | 27 | 222 | 672 | 672 |
| Lm | 5 | 38 | 44.0 | 977 | 562 | 1.391 | 22 | 623 | 0.0 | 0.0 |

### Engagement is maintained at larger action-reward separation

To test whether e-Seesaw use was restricted to short action–reward separations, we examined an additional subset of deployments in which the distance between the interaction site and reward module was increased to 100 cm (Fig. 1H). Across three families and 33 sessions, animals continued to engage with the device, producing a median of 38 trigger activations within the first 20 min after the first press (IQR, 28–41; range, 13–57), (Table S1). This level of early-session engagement was within the range observed in the core 20-cm deployment, where sessions showed a median of 37 activations in the same time window (IQR, 23–54). Across repeated 100-cm sessions, engagement did not decline, with two families showing nominally positive slopes across sessions, supporting the feasibility of larger spatial dissociations between action and reward.

## Discussion

We developed the e-Seesaw system as a home-eclosure-compatible system in which reward delivery is spatially dissociated from the site of action. This flexible, low cost and modular design makes it possible to study situations in which one individual’s action can affect reward availability without requiring experimenter-imposed dyads, repeated handling, or enclosure modifications. Crucially, animals can interact with the system within their routine group environment, participation remains voluntary, and the opportunity for partner choice is preserved rather than being structured by the experimenter. Furthermore, removing the experimenter from the testing situation prevents a potential confound that monkeys may interpret the experimenter as a social agent participating in the exchange itself, rather than as a neutral observer (*42, 55*). The present study therefore focused on deployment feasibility and usage dynamics under home-enclosure conditions, a necessary first step before extending the system to paradigms that assign actions and outcomes to specific individuals in social space.

A central outcome of the proof-of-concept deployment is that group-housed marmosets engaged reliably with the device across families and sessions, with rare zero-trigger sessions (Fig. 1B), demonstrating feasibility under routine social housing where devices must sustain voluntary use alongside daily husbandry (*56*). Engagement was not uniformly distributed over time, but strongly front-loaded within sessions (Fig. 1E) and, when animals engaged, inter-trigger gaps were short (Fig. 1F), consistent with sustained bouts of use. This pattern suggests that the earliest portion of a session is the most informative about reward-directed engagement, whereas later triggers are more ambiguous because reward depletion did not define session termination, allowing presses to occur even after reward was no longer available. Consistent with this interpretation, total session duration explained only a small fraction of the variance in trigger counts, indicating that longer recording windows do not proportionally increase interaction (Fig. 1G). Together, these patterns argue against treating engagement as a simple time-normalized rate and motivate metrics that reflect structured, state-dependent behavior in unconstrained settings (*57, 58*).

A second important outcome concerns the spatial separation of action and reward. This feature was built to capture a core feature of paradigms such as the PCT and group service task (*15, 24, 29*), in which one individual’s action determines reward access for others. The present dataset addresses a practical concern of this design, namely whether animals would continue to engage when the reward module is moved farther from the trigger. In 100-cm deployments across three families and 33 sessions, animals produced a median of 38 trigger activations similar to the median of 37 activations observed in the core 20-cm deployment (Fig. 1H). This demonstrates that engagement was not restricted to short action–reward separations, and is fully consistent with marmosets’ prosocial tendencies in traditional paradigms (*59, 60*).

Two limitations define the current scope of inference. First, because this study does not identify individual actors or measure consumption, claims about actor–recipient relationships, monopolization, or equity of access are not yet supported. These same limitations reflect a broader practical bottleneck in social housing, where access to baseline social patterns often require extensive manual observation. Integrating individual attribution (e.g., RFID or video-based identification) will allow e-Seesaw logs to contribute to automated baselining around reward delivery, complementing existing approaches to social behavior quantification (*61*–*63*), and will also provide a necessary foundation for future studies linking social behavior in group settings to individual-level neural activity. Second, because engagement is strongly front-loaded, later portions of long sessions contribute limited incremental information about participation under the current deployment regime. This motivates early-window metrics but also points to future experiments that explicitly test how schedules, effort, novelty, or social context shape engagement dynamics.

In sum, the e-Seesaw provides an open, enclosure-compatible method for wireless, spatially displaced reward delivery in marmosets, with robust engagement across families and largely stable participation across sessions. By combining autonomous operation with flexible geometry between action and reward sites, the platform addresses a methodological bottleneck for studying reward access in social space. The modular design should also be readily adaptable to alternative trigger mechanisms, such as buttons or other actuators, which could extend the framework to different task structures and to larger-bodied species. With future integration of individual identification and complementary sensing, the same framework could further reduce the burden of establishing social baselines in captive groups and support more direct tests of reward allocation under routine housing conditions.

## Methods

### Animals

We studied 7 captive family groups of common marmosets (*Callithrix jacchus*), each consisting of 4–5 members. Families were maintained in heated indoor enclosures (2.7*1.8*2.4 m) with outdoor access when weather conditions permit (3.2*1.8*2.4 m). Family groups are housed in proximity with visual occlusion between groups but a shared acoustic environment. All animals had ad libitum access to water, were fed at least three times daily in the morning with a vitamin enriched porridge, around midday with fresh vegetables and a variety of proteins such as beans and in the late afternoon with insects and arabic gum. All procedures were approved by Kantonales Veterinäramt Zürich, license number ZH211/2022.

### Hardware and software architecture

The e-Seesaw comprises of three functional elements: (i) a server module that receives triggers via bluetooth, controls reward delivery, and logs data, (ii) a client module embedded in the mechanical seesaw apparatus that detects activation and transmits triggers wirelessly, and (iii) the mechanical seesaw apparatus that couples animal interaction to the trigger signal (Fig. S1-3).

The server module is built around a Raspberry Pi Zero that controls a peristaltic pump (Adafruit Peristaltic Liquid Pump, 5–6 V) to dispense liquid reward from a 50 mL reservoir. The module includes a main power switch (battery bank powered) and a secondary manual-prime switch that directly activates the pump for priming and tubing fill. A status LED indicates server power state. Wiring schematics are provided in Fig. S1.

The client module is built around an Arduino Micro coupled to an HC-05 Bluetooth module for wireless communication with the server. Seesaw activation is detected by a simple contact mechanism at the hinge: two stainless-steel fasteners close an electrical circuit when brought into contact by seesaw motion, generating a trigger event. The client includes a power switch (battery bank powered) and two status LEDs controlled by the Arduino: one indicates power state and the other indicates successful server pairing. Wiring schematics are provided in Fig. S2.

The mechanical seesaw is a fully 3D-printed assembly designed to provide a robust, enclosure-mountable interface (Fig. S3). It consists of three main printed parts: (i) an outer and inner grid fixture (Fig. S3D–E) that provides the anchor supporting the seesaw mechanism on the grid of the enclosure (ii) a horizontal bar (Fig. S3A) that connects via two hinges to an outer plate (Fig. S3C) that extends via two arms into the inside of the enclosure where (iii) the monkeys are provided with a sitting plate (Fig. S3B). The outer plate provides anchor points to mount the client module (Fig. S3B), including routing points for the two lead wires connected to the hinge contact.

On power-up, the client enters pairing mode and the server attempts to connect automatically. Pairing on the Raspberry Pi is executed via a scheduled startup routine (crontab calling a bash wrapper and a Python script). If pairing fails, the user restarts both modules. Once paired, the system is ready for operation.

Communication between modules is managed by a custom Python program on the Raspberry Pi. The server establishes a Bluetooth handshake with the Arduino client, listens for trigger messages, and on each trigger actuates the peristaltic pump to deliver a pre-specified volume of reward. The program logs trigger events with timestamps and associated delivery parameters and saves outputs in CSV format for downstream analysis. Data is automatically backed up to a local server over Wi-Fi.

We selected the Raspberry Pi to simplify integration of additional peripherals and sensors through standard interfaces (GPIO, SPI, I2C). Both the Raspberry Pi server and Arduino-based client can be operated from portable power banks, allowing flexible placement within the animal facility while avoiding direct reliance on mains power near the enclosure. This low-voltage, battery-based configuration reduces electrical risk in the event of accidental contact or cable damage and improves portability across deployments. Furthermore, for replication across devices, new units can be provisioned by duplicating the configured SD card image from an existing server module, enabling consistent software setup across deployments.

### Apparatus design, construction and implementation

Custom housings for the server and client modules were 3D-designed and printed in-house, and mounted to the enclosure’s wire grid such that electronics and fluidics remained outside the cage while the seesaw interface extended inside for animal interaction. STL/CAD files, a complete bill of materials, wiring diagrams (Fig. S1-3), assembly (Fig. S4) and setup instructions are provided in the Supplementary Materials and the project repository (GitHub: jorgecabreira/e-seesaw).

### Prior familiarization with seesaw mechanism

All groups included in the present dataset had prior experience with a functionally similar seesaw apparatus as part of parallel husbandry and experimental procedures in the colony. The initial training to learn how to operate this seesaw took place between July and December 2023, at least one year before the e-Seesaw deployments analyzed here, and data from that phase are not included in the present study. The training was conducted in sessions of 20 minutes per family. The training had three phases. First the bottle containing the liquid reward was installed very close to the seesaw platform, so that the animal triggering the apparatus could lick the reward themselves without moving. In phase two the bottle with the reward was further away so that the animal could still reach it when stretching themselves from the seesaw platform and in the last phase the bottle was at the other end of the apparatus not reachable for the animal triggering the mechanism. To pass to the next stage of training every animal in each group had to trigger the seesaw and lick from the reward bottle within 3 seconds at least 10 times in 5 separate sessions.

### e-Sessaw deployment

The e-Seesaw was installed permanently in each home enclosure (starting around 07:00 - 08:00 in the morning, see Table S2) and manually turned on by caretakers around 8:00 am during their routine feeding round, dispensing 100–200ml of morning porridge (rice flakes - Alnatura, Arabic gum, and water). During initial sessions, the server and client modules were positioned in close proximity, such that triggering the seesaw delivered immediate self-reward. For the core feasibility dataset, the reward module was positioned 20 cm from the seesaw interaction site. Additional larger-distance deployments were conducted in a subset of families and are reported separately as supplementary feasibility observations (Table S1).

## Acknowledgements

We would like to thank Hidir Sengül and Eileen Osswald for their support in keeping all the animals healthy and especially for incorporating the e-Seesaw during their caretaking routine. We thank Laura Andres and Henry Broadbent for developing the apparatus that was used in the initial training phase, and for training the animals to use the seesaw mechanism. We thank Eileen Osswald and Carolyn Sanicanin for their help during the deployment phase. We are grateful to Andreas Hess for his input in developing the 3d models and 3d-printing the e-Seesaw parts.

## Availability of data and materials

The datasets and analysis code required to reproduce the results will be deposited in a Zenodo repository. During peer review, these materials will be made available to editors and reviewers through a private Zenodo preview link. Upon publication, the repository will be made publicly available and citable through a Zenodo DOI.

## Authors Contributions

**J.C.M**.: Conceptualization, Methodology, Software, Investigation, Data curation, Formal analysis, Visualization, Writing – original draft, Writing – review and editing. **J.M.B**.: Funding Acquisition, Resources, Supervision, Writing – review and editing. **R.K.B**.: Conceptualization, Methodology, Investigation, Visualization, Project administration, Writing – original draft, Writing – review and editing.

All authors gave final approval for publication.

