## Supplemental Figure S1 to S4, Table S1 to S2 for "An Automated Wireless Seesaw System Enabling Spatial Separation of Action and Reward in Group-Housed Marmosets"

Content:
Figure S1 to S4
Table S1 to S2

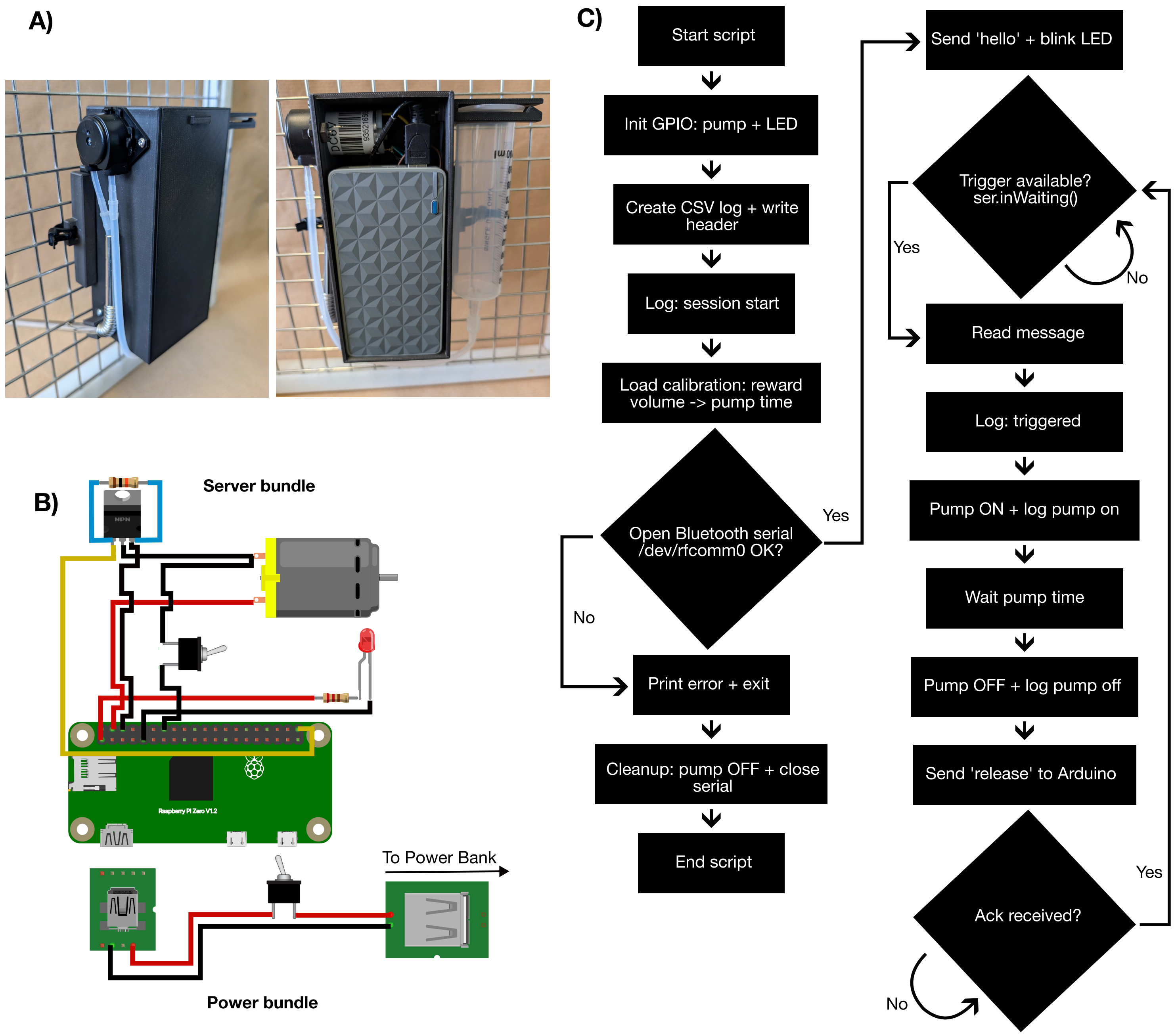


**Figure S1. e-Seesaw server module: hardware, wiring, and control logic.**
(A) Photograph of the assembled server module housed in the protective enclosure. The server contains the Raspberry Pi Zero, pump driver circuitry, and external connectors for power and the peristaltic pump. (B) Wiring schematic of the server electronics. The Raspberry Pi controls the peristaltic pump via a GPIO-driven output (MOSFET switch) and provides a status LED output; the peristaltic pump draws power from the external battery supply. (C) High-level workflow of the Raspberry Pi control program. On startup, the server initializes GPIO and logging, loads the pump calibration (target reward volume mapped to pump-on time), and establishes a Bluetooth serial connection to the client. During runtime, each trigger received from the client is time-stamped and logged, the pump is activated for the calibrated duration, pump ON/OFF events are logged, and a release message is sent back to the client to acknowledge completion and re-arm the trigger.

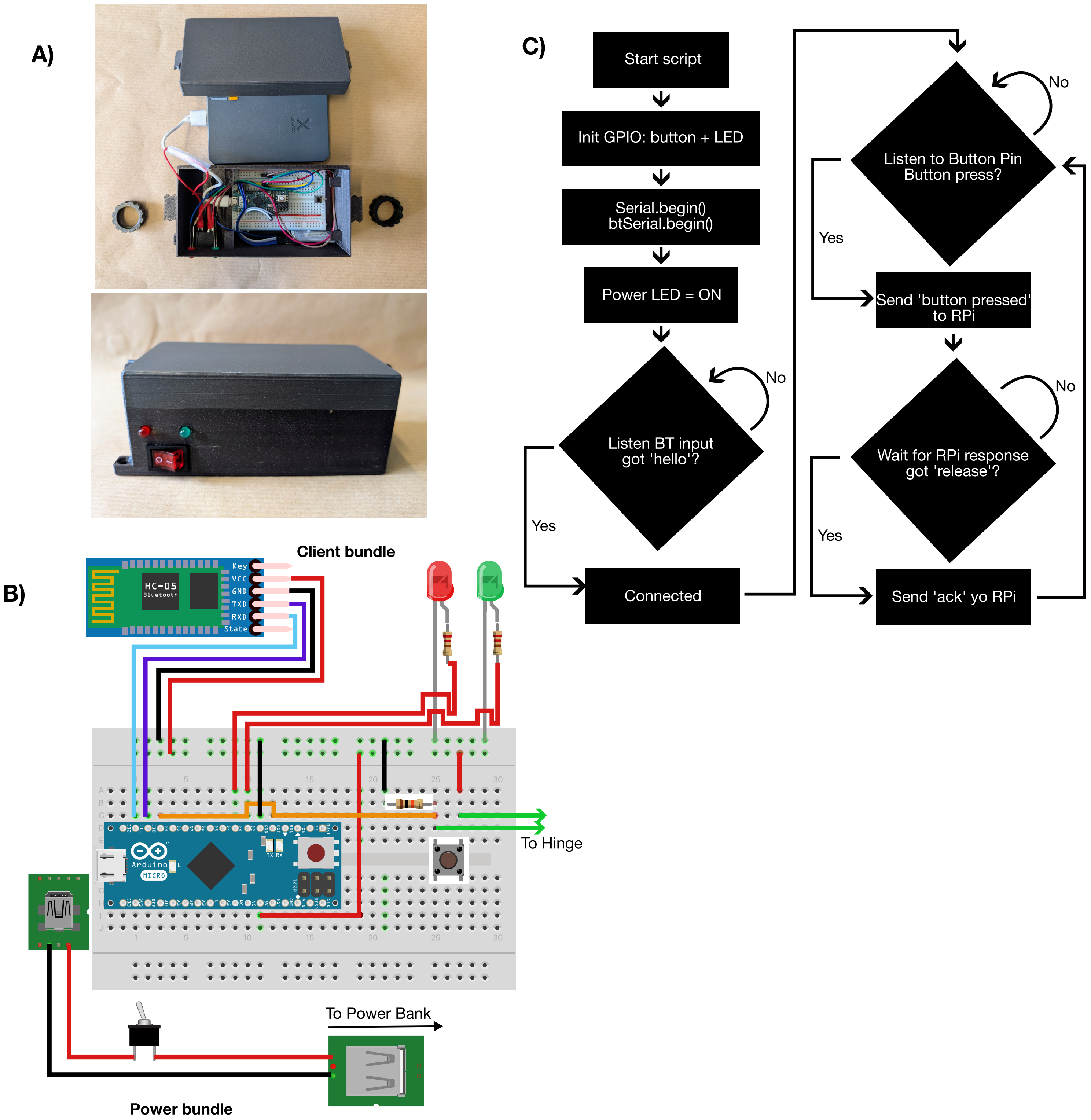


**Figure S2. e-Seesaw client module: hardware, wiring, and communication logic.**
(A) Photograph of the assembled client module mounted on the seesaw. The client houses the Arduino Micro and HC-05 Bluetooth module and interfaces with the hinge contact used to detect seesaw activation. (B) Wiring schematic of the client electronics. The hinge contact is read as a digital input by the Arduino, and two status LEDs indicate client power state and successful Bluetooth pairing with the server. (C) High-level workflow of the client firmware. After startup, the client initializes I/O and Bluetooth serial communication, then waits for a handshake message ("hello") from the Raspberry Pi server to confirm a valid connection. Once connected, seesaw activation triggers transmission of a "Button pressed" message to the server. The client then waits for a "released" message indicating reward delivery completion, replies with an "ack" acknowledgment, and enforces a button-release hold-off (debounce) before accepting the next trigger.

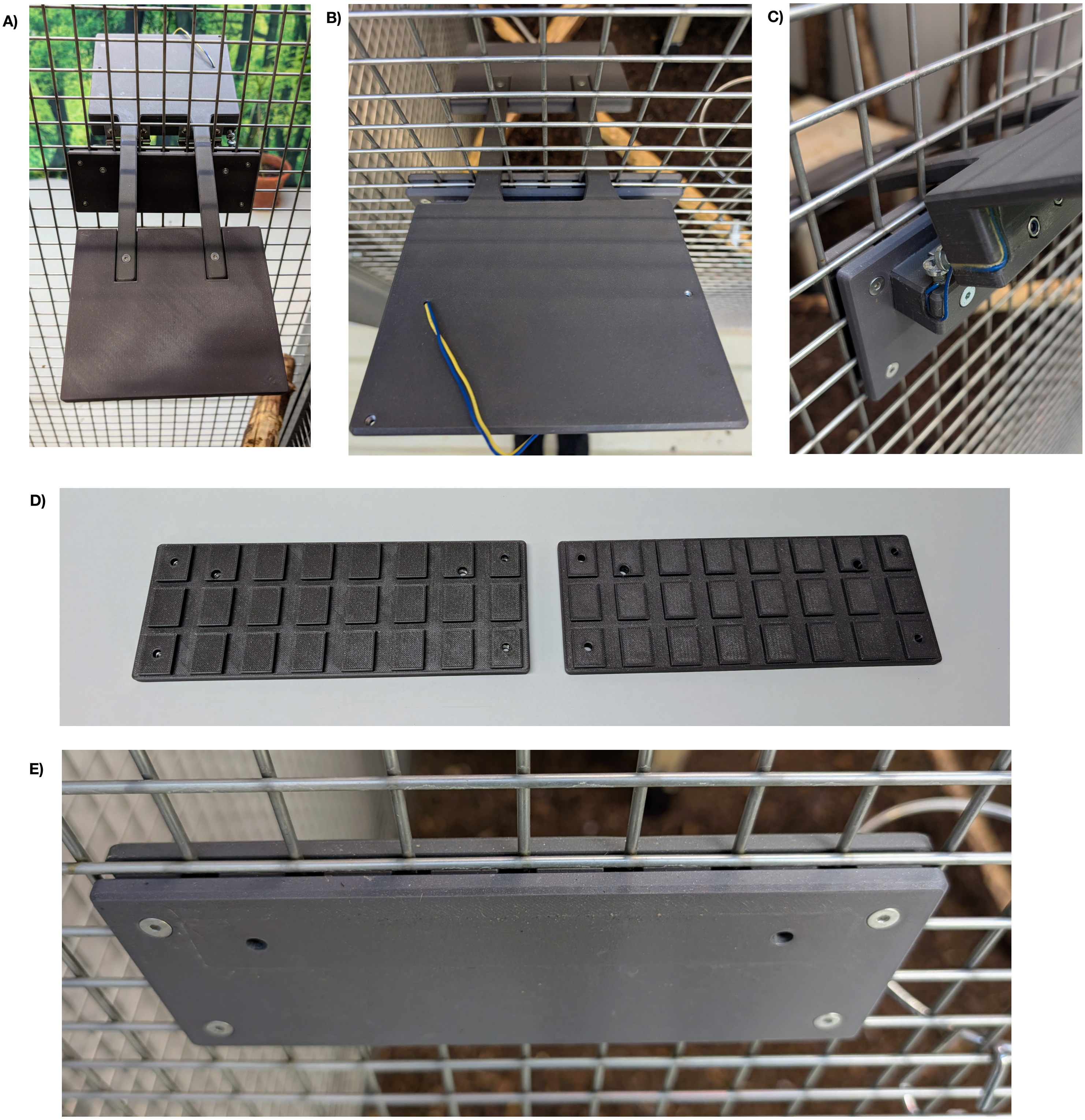


**Figure S3. Mechanical seesaw components and enclosure mounting.**
(A) View from inside the cage. The inner platform is attached with screws to the two arms of the outer platform. This inner platform serves as the primary animal interaction surface used to actuate the seesaw. (B) View from outside the cage. The outer platform extends inward through the cage wall to support the inner platform and also contains the mounting points for the client module, as well as the routing path for the hinge contact leads. (C) Close-up of the cable routing. The cables are positioned so that they make contact with the screws, closing the circuit when the seesaw is touched or triggered. (D) Image of the grid-anchor panels. (E) Image of the grid-anchor panels mounted onto the cage. The grid-anchor component clamps onto the enclosure wire mesh and provides structural support for the hinge axis, ensuring stable positioning of the seesaw assembly.

| | # | size | type | ISO | nut | type |
|---|---|---|---|---|---|---|
| **A** | 12 | M4 x 22 mm | Hexagon socket button head screw with collar (stainless steel A2) | ISO 7380-2 | M4 | Nyloc nut ISO 10511 (stainless steel A2) |
| **B** | 4 | M4 x 20 mm | Hexagon socket countersunk head cap screw (stainless steel A2) | ISO 10642 | M4 | Nyloc nut ISO 10511 (stainless steel A2) |
| **C** | 2 | M5 x 45 mm | Hexagon socket countersunk head cap screws (steel FKL 8.8) | ISO 10642 | M5 | Nyloc nut ISO 10511 (stainless steel A2) |
| **D** | 2 | M4 x 12 mm | Hexagon socket countersunk head cap screw (stainless steel A2) | ISO 10642 | M4 | Nyloc nut ISO 10511 (stainless steel A2) |
| **E** | 2 | M4 x 20 mm | Hexagon socket countersunk head cap screw (stainless steel A2) | ISO 10642 | M4 | Nyloc nut ISO 10511 (stainless steel A2) |
| **F** | 2 | M5 x 16 mm | Hexagon socket head cap screw (steel galvanised) | ISO 4762 | - | - |

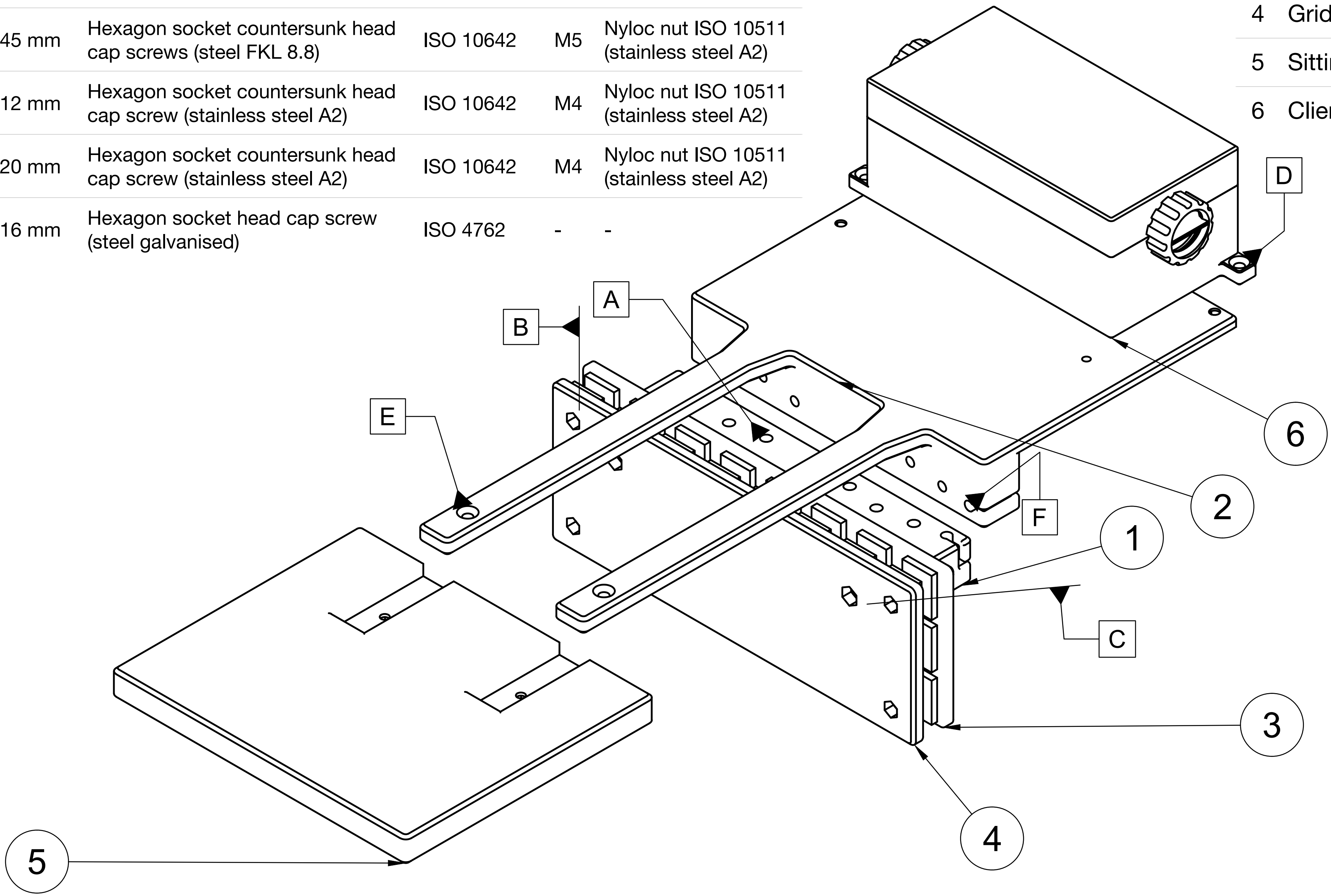


| | |
|---|---|
| 1 | Horizontal bar outside |
| 2 | Plate outside |
| 3 | Grid fixture outside |
| 4 | Grid fixture inside |
| 5 | Sitting plate inside |
| 6 | Client box outside |

**Supplementary Figure S3.** Assembly guide for the seesaw mechanism of the mechanical seesaw. The horizontal bar outside (1) and the plate outside (2) are connected via two hinges (that are fixed with scews A). Two lead wires are glued into the routing points and connected to the two hexagon socket head cap screws, which are inserted into the holes (F). The grid fixture outside and inside is mounted onto the grid (with screws B), and then the horizontal bar outside is fixed onto the grid anchor (with screws C). Lastly, the sitting plate is screwed onto the extensions of the outside plate (with screws E). The client box can be mounted onto the outside plate (with screws D).

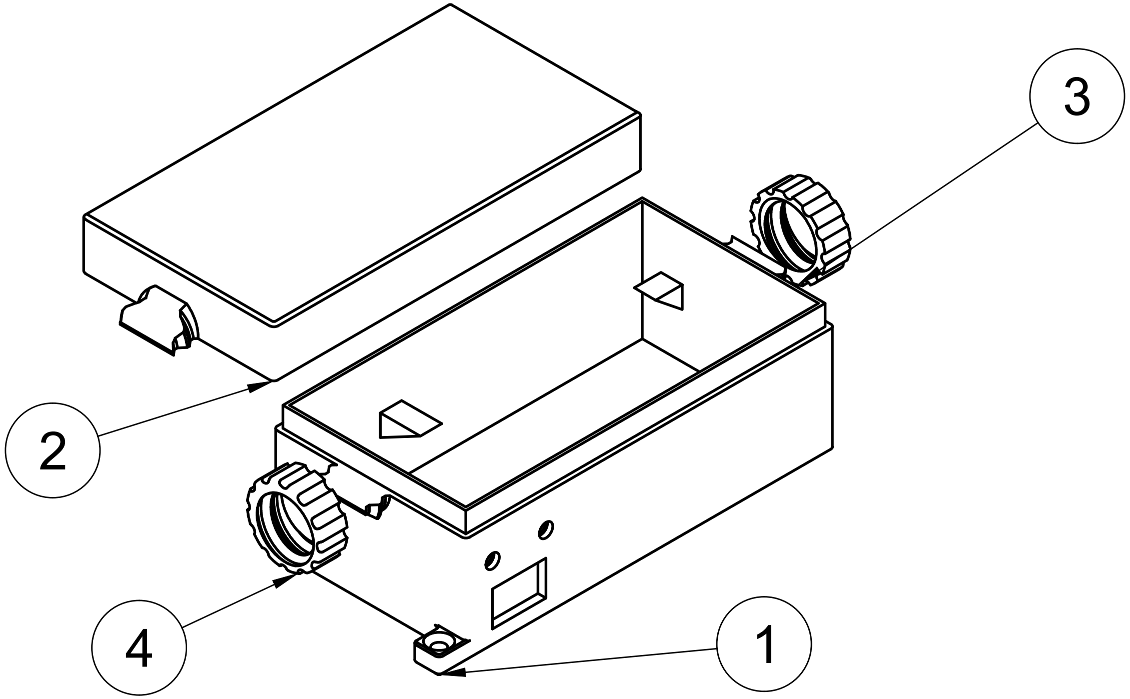


| | |
|---|---|
| 1 | Client box bottom |
| 2 | Client box lid |
| 3 | Client box closing screw right |
| 4 | Client box closing screw left |

**Supplementary Figure SXX.** Assembly guide for the client box. The client box has a bottom part with holes for the power switch, the LEDs, and the lead cables, as well as triangular platforms to hold the battery in place. It is closed with a lid that is screwed on from the outside with two closing screws.

| | # | size | type | ISO | nut | type |
|---|---|---|---|---|---|---|
| **A, B, C** | 6 | M3 x 12 mm | Hexagon socket button head screw (stainless steel A2) | ISO 7380-1 | M3 | Nyloc nut ISO 10511 (stainless steel A2) |
| **D** | 4 | M2.5 x 10 mm | Torx socket countersunk head cap screw (stainless steel A2) | ISO 14581 | – | – |

| | |
|---|---|
| 1 | Server box |
| 2 | Server box lid |
| 3 | Server box holder |
| 4 | Straw holder |
| 5 | Syringe holder |

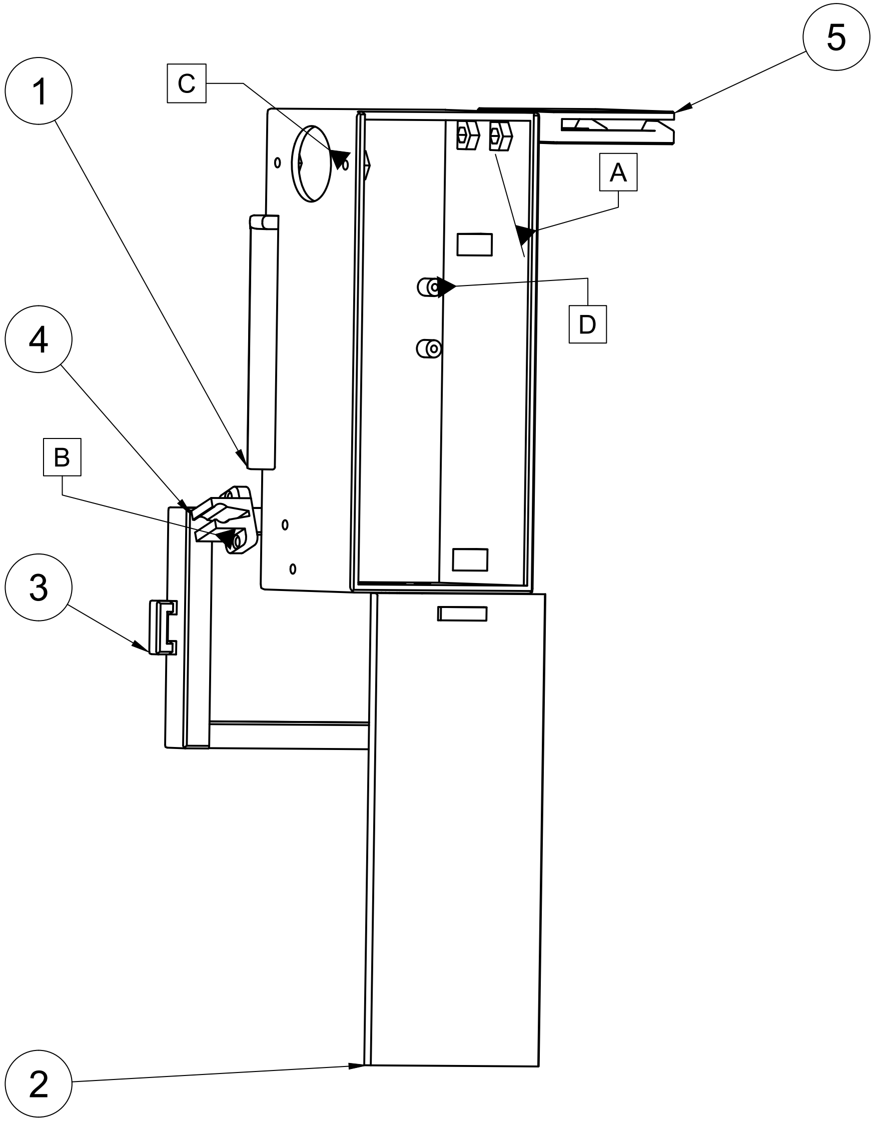


**Supplementary Figure SXX.** Assembly guide for the server box. The server box has a bottom part with holes for the power switch, the manual switch for the pump, the LEDs, the pump, and the straw holder. The straw holder and the syringe holder are screwed on with screws A and B. There are little screew holes to mount the Raspberry Pi with screws D as well as triangular platforms to hold the battery in place. The box is closed with a sliding lid. The server box is mounted onto the grid with a holder that can be fixed with zipties.

| family | n_sessions | median_presses | slope_per_session | slope_ci_lo | slope_ci_hi | relative_slope | pearson_r | p_slope | p_slope_fdr |
|---|---|---|---|---|---|---|---|---|---|
| **Lm** | 12 | 36.5 | 0.26 | -2.11 | 2.64 | 0.01 | 0.08 | 0.81 | 0.81 |
| **Gr** | 10 | 40 | 1.86 | 0.02 | 3.69 | 0.05 | 0.64 | 0.05 | 0.07 |
| **Gp** | 11 | 29 | 1.76 | 0.07 | 3.44 | 0.06 | 0.62 | 0.04 | 0.07 |

**Table S1.** Summary of engagement for deployments at 100 cm distance between the trigger and the reward module. For each family, the table reports, the group size, the number of sessions analyzed, the median number of triggers within the first 20 minutes of a session, the estimated linear slope of early-session triggers per session (with 95% confidence interval), the relative slope (slope normalized by the family median), the Pearson correlation coefficient, the slope p-value and FDR-corrected p-value.

| family | distance between trigger and reward | number of sessions | median session duration ± SE | start data collection | end data collection | Group composition |
|---|---|---|---|---|---|---|
| Gp | 20 cm | 9 | 96.9 ± 19.8 | 2025-02-19 | 2025-03-20 | fb, mb, fh, mh |
| Gp | 100 cm | 11 | 11.4 ± 16.1 | 2026-04-08 | 2026-04-21 | fb, mb, fh, mh |
| Gr | 20 cm | 21 | 90.9 ± 26.1 | 2025-03-21 | 2025-05-08 | fb, mb, fh, mh |
| Gr | 100 cm | 10 | 86.9 ± 10.6 | 2026-03-21 | 2026-04-28 | fb, mb, fh, mh |
| Jj | 20 cm | 11 | 202.1 ± 47.1 | 2025-03-03 | 2025-03-17 | fb, mb, fh, mh |
| Jn | 20 cm | 6 | 80.5 ± 10 | 2025-03-03 | 2025-03-10 | fb, mb, 2 mh, fh |
| Lm | 20 cm | 38 | 101.1 ± 8.2 | 2025-03-26 | 2025-06-06 | fb, mb, 2 fh, mh |
| Lm | 100 cm | 12 | 91 ± 15.7 | 2026-04-15 | 2026-04-29 | fb, mb, 2 fh, mh |
| Mj | 20 cm | 10 | 127.5 ± 50.5 | 2025-03-11 | 2025-03-25 | fb, mb, 2 fh |
| Ml | 20 cm | 6 | 87.9 ± 49.8 | 2025-02-21 | 2025-02-28 | fb, mb, 2 fh, mh |

**Table S2.** Overview of subjects participating in e-seesaw deployments. Family groups and their group compositions at the time of data collection. Mean session duration  fb = female breeder, mb = male breeder, fh = female helper, mb = male helper
